# Characterizing and predicting cyanobacterial blooms in an 8-year amplicon sequencing time-course

**DOI:** 10.1101/058289

**Authors:** Nicolas Tromas, Nathalie Fortin, Larbi Bedrani, Yves Terrat, Pedro Cardoso, David Bird, Charles W. Greer, B. Jesse Shapiro

**Author notes:** Corresponding authors: B. Jesse Shapiro.; Nicolas Tromas.

## Abstract

Cyanobacterial blooms occur in lakes worldwide, producing toxins that pose a serious public health threat. Eutrophication caused by human activities and warmer temperatures both contribute to blooms, but it is still difficult to predict precisely when and where blooms will occur. One reason that prediction is so difficult is that blooms can be caused by different species or genera of cyanobacteria, which may interact with other bacteria and respond to a variety of environmental cues. Here we used a deep 16S amplicon sequencing approach to profile the bacterial community in eutrophic Lake Champlain over time, to characterize the composition and repeatability of cyanobacterial blooms, and to determine the potential for blooms to be predicted based on time-course sequence data. Our analysis, based on 135 samples between 2006 and 2013, spans multiple bloom events. We found that bloom events significantly alter the bacterial community without reducing overall diversity, suggesting that a distinct microbial community – including non-cyanobacteria – prospers during the bloom. We also observed that the community changes cyclically over the course of a year, with a repeatable pattern from year to year. This suggests that, in principle, bloom events are predictable. We used probabilistic assemblages of OTUs to characterize the bloom-associated community, and to classify samples into bloom or non-bloom categories, achieving up to 92% classification accuracy (86% after excluding cyanobacterial sequences). Finally, using symbolic regression, we were able to predict the start date of a bloom with 78-92% accuracy (depending on the data used for model training), and found that sequence data was a better predictor than environmental variables.

## Introduction

Cyanobacterial blooms occur in freshwaters systems around the world, and are both a nuisance and a public health threat (Zingone and Enevoldsen, 2000; Paerl *et al*., 2013). These blooms are defined by a massive accumulation of cyanobacterial biomass, formed through growth, migration, and physical–chemical forces (Paerl, 1996). In temperate eutrophic lakes, blooms tend to occur annually, specifically during the summer when water temperatures are warmer (Kanoshina *et al*., 2003, Havens, 2008). The frequency and intensity of these blooms is increasing over time (Johnson *et al*., 2010; Posch *et al*., 2012), likely due to increased eutrophication, climate change, and increased nutrient input from human activities (O’Neil *et al*., 2012; Winder, 2012).

Attempts have been made to predict blooms using hydrodynamic-ecosystem models (Allen, 2008, Qing *et al*., 2014), artificial neural networks models (Maier *et al*., 2000; 2001), or statistical models such as on linear regression (Dillion and Rigler, 1974, Onderka, 2007). Nevertheless, these models have been limited in their ability to accurately predict cyanobacterial dynamics (Downing *et al*., 2001; Taranu *et al*., 2012), perhaps because they mainly used abiotic factors (*e.g.* temperature, pH, nutrients, etc.) to predict blooms, while largely ignoring biotic factors (Recknagel *et al*., 1997; Downing *et al*., 2001; Oh *et al*., 2007). It is known that cyanobacteria interact with their biotic environment in a variety of ways, ranging from predator-prey interactions to mutualistic interactions (Rashidan and Bird, 2001; Eiler and Bertilsson, 2004; Berg *et al*., 2008; Li *et al*., 2012; Mou *et al*., 2013; Louati *et al*., 2015; Woodhouse *et al.* 2016). Biotic factors, such as the composition of the surrounding bacterial community, could therefore help refine bloom prediction. Previous studies have predicted the distribution of other bacteria based on community structure (Larsen *et al* 2012; Kuang *et al*., 2016) but to our knowledge this has not been attempted to predict freshwater cyanobacterial blooms. Prediction based on biotic factors is attractive because the composition of the microbial community can be thoroughly measured through culture-independent, high-throughput sequencing, whereas it is not always clear which are the relevant (or most predictive) abiotic factors that should be measured. Moreover, the microbial community composition may contain information about both measured and unmeasured abiotic variables, insofar as these variables impact the community.

For bloom prediction based on biotic factors to be successful, there must be some degree of repeatability in the changes to lake bacterial community composition that precede blooms. Several studies have shown that many aquatic microbial communities are temporally dynamic (Pernthaler *et al*., 1998; Hofle *et al*., 1999; Lindstrom *et al*., 2000; Crump et al., 2003; Kent *et al.*, 2004; Shade *et al*., 2007; Kara *et al*., 2013; Fuhrman *et al*., 2015), often with repeatable patterns of community structure (Fuhrman *et al*., 2006; Fuhrman *et al*., 2015). Recent studies have tracked the dynamics of microbial communities in bloom-impacted lakes using culture-independent sequencing methods (Eiler *et al*., 2012; Li *et al*., 2015; Woodhouse *et al*., 2016). Li *et al*. (2015) found that a bloom-impacted lake returned to its initial community composition after a period of one year. However, all these studies were carried out over one year or less, making it difficult to generalize the results and make robust predictions. As highlighted by Fuhrman *et al*., (2015) data should be collected over several consecutive years to assess the repeatability of bacterial community dynamics, and to assess if community structure follows a predictable pattern, and over what time scales.

Blooms can be operationally defined in numerous ways. A classic definition is simply when algal biomass is high enough to be visible (Reynolds and Walsby, 1975). Other bloom definitions rely on chlorophyll concentrations (≥20 μg/L), or dominance of cyanobacteria (>50%) over other phytoplankton (Molot *et al*., 2014). An attractive alternative is to view cyanobacterial blooms as a biological disturbance, measurable by their impact on the surrounding microbial community (Shade *et al*., 2012). Blooms can have a major impact on the microbial community through both direct (e.g. microbe-microbe interactions) and indirect effects (e.g. changes to lake chemistry). For example, blooms can reduce carbon dioxide concentrations, increase pH, and alter the distribution of biomass across the length and depth of a lake (Verspagen *et al.,* 2014; Sandrini *et al*., 2016). Such bloom-induced changes in water chemistry could then impact the structure and diversity of microbial communities (Bouvy *et al*., 2001; Eiler and Bertilsson, 2004; Bagatini *et al*., 2014; Li *et al*., 2015, Woodhouse *et al*., 2016). For example, as cyanobacteria decompose, they release metabolites that can be utilized by other taxa, such as Cytophagaceae (Rashidan and Bird, 2001; O’Neil *et al*., 2012), which we therefore expect to be observed in association with blooms. Positive associations have been observed between the genus *Phenylobacterium* or members of the order Rhizobiales with the cyanobacterial genus *Microcystis* (Louati *et al*., 2015). However, the reasons for these interactions, as well as their repeatability (over time) and generality (across different lakes) remain unknown.

Here, we present an eight-year time-course study of the bacterial community structure of a large eutrophic North American lake, Lake Champlain, where cyanobacterial blooms are observed nearly every summer. Samples were collected from 2006 to 2013 and analyzed using high-throughput 16S amplicon sequencing. We tracked the bacterial community composition in 135 time-course samples to determine how the community varies over time and how it is impacted by blooms. Considering blooms as a disturbance to the surrounding microbial community (Shade *et al.* 2012), we defined bloom events as a relative abundance of cyanobacteria above which community diversity begins to decline. Blooms are characterized both by a dominance of cyanobacteria, but also a characteristic surrounding bacterial community. We show that the community composition does not vary considerably from year to year, but does vary within a year, on time scales of days to months. As a result, community dynamics are largely repeatable from year to year, and are in principle predictable. Finally, exploiting the repeatable dynamics of the lake community, we showed that bloom events can be predicted several weeks in advance based on the microbial community composition, with slightly greater accuracy than predictions based on abiotic factors.

## Materials and Methods

### Sampling

A total of 150 water samples were collected from the photic zone (0-1 meter depth) of Missisquoi Bay, Lake Champlain, Quebec, Canada (45°02'45''N, 73°07'58''W). Between 12 and 27 (median 17) samples were collected each year, from 2006 to 2013, between April and November of each year. Samples were taken from both littoral (78 samples) and pelagic (72 samples) zones (Supplementary Methods). Between 50 and 250 ml of lake water was filtered depending on the density of the planktonic biomass using 0.2-μm hydrophilic polyethersulfone membranes (Millipore). Physico-chemical measurements, as described in Fortin *et al*. (2015), were also taken during most sampling events (Supplementary File: File_S1_Environmental_Table.txt). These environmental data included water temperature, average air temperature over one week, cumulative precipitation over one week, microcystin toxin concentration, total and dissolved nutrients (phosphorus and nitrogen). Details of the sampling protocol are described in Supplementary Methods.

### DNA extraction, purification and sequencing

DNA was extracted from frozen filters by a combination of enzymatic lysis and phenol-chloroform purification as described by Fortin *et al*. (2010). Each DNA sample was resuspended in 250 μl of TE (Tris-Cl, 10 mM; EDTA, 1 mM; pH 8) and quantified with the PicoGreen^®^ dsDNA quantitation assay (Invitrogen). DNA libraries for paired-end Illumina sequencing were prepared using a two-step 16S rRNA gene amplicon PCR as described in Preheim *et al*. (2013). We amplified the V4 region, then confirmed the library size by agarose gels and quantified DNA with a Qubit v.2.0 fluorometer (Life Technologies). Libraries were pooled and denatured as described in the Illumina protocol. We performed two sequencing runs using MiSeq reagent Kit V2 (Illumina) on a MiSeq instrument (Illumina). Each run included negative controls and two mock communities composed of 16S rRNA clones libraries from other lake samples (Preheim *et al*., 2013). Details of the library preparation protocol are described in Supplementary Methods.

### Sequence analysis and OTU picking

Sequences were processed with the default parameters of the SmileTrain pipeline (https://github.com/almlab/SmileTrain/wiki/; Supplementary Methods) that combined reads quality filtering, chimera filtering, paired-end joining and, de-replication using USEARCH (version 7.0.1090, http://www.drive5.com/usearch/) (Edgar, 2010), Mothur (version 1.33.3) (Schloss *et al*., 2009), Biopython (version 2.7) and custom scripts. SmileTrain also incorporates a *de novo* distribution-based clustering: dbOTUcaller algorithm (Preheim *et al*., 2013) (https://github.com/spacocha/dbOTUcaller, version 2.0), which was performed to cluster sequences into Operational Taxonomic Units (OTUs) by taking into account the sequence distribution across samples. The OTU table generated was then filtered using filter_otus_from_otu_table.py QIIME scripts (Caporaso *et al*., 2010) (version 1.8, http://qiime.org) to remove OTUs observed less than 10 times, minimizing false-positive OTUs (Table S1). Fifteen samples with less than 1000 sequences were removed from the OTU table using filter_samples_from_otu_table.py QIIME script, yielding a final dataset of 135 samples. Taxonomy was assigned post-clustering using a two different approaches: (i) the latest 97% reference OTU collection of the GreenGenes database (release 13_8, August 2013, ftp://greengenes.microbio.me/greengenes_release/gg_13_5/gg_13_8_otus.tar.gz; http://greengenes.lbl.gov), using assign_taxonomy.py QIIME script (default parameters), and (ii) a combination of GreenGenes and a freshwater-specific database (Freshwater database 2016 August 18 release; Newton *et al*., 2011), using the TaxAss method (https://github.com/McMahonLab/TaxAss, access date: September 13th 2016). Taxonomy information was then added to the OTU table using the biom add-metadata scripts (http://biom-format.org). We removed OTUs that were not prokaryotes but still present in the database (Cryptophyta, Streptophyta, Chlorophyta and Stramenopiles orders). A total of 7,321,195 sequences were obtained from our 135 lake samples, ranging from 1,392 to 218,387 reads per sample, with a median of 47,072. This dataset was clustered into 4061 OTUs. Of these OTUs, 4053 were observed in littoral samples and 4042 in pelagic samples, with 4034 in common to both sites, 19 unique to littoral and 8 to pelagic.

To evaluate the quality of the SmileTrain OTU picking pipeline used and estimate the potential false positive OTUs generated by the approach used, we compared the number and identity of OTUs obtained for two different mock communities that were generated from plasmids containing 16S rRNA sequences from a clone library as described on Preheim *et al.* (2013). SmileTrain (using the dbOTUcaller algorithm) recovered 100% of the expected OTUs in the mock community, *i.e* we found a perfect match between 16S sequences from the library and the OTU representative sequences generated post-clustering. However we also found some false positives (Table S1). We removed OTUs represented by fewer than 10 sequences in total to minimize false positives using filter_otus_from_otu_table.py QIIME script. (Table S1). After this filtering, we still recovered 97% for Mock10 and 100% for Mock11. Details of the post-sequencing computational pipeline are described in Supplementary Methods, and R scripts (for analyses described here and below) are in Supplementary File 2 (File_S2_R_scripts.txt).

### Diversity analysis

To calculate the alpha diversity, indexes known for their robustness to sequencing depth variation were used: Shannon diversity (Shannon and Weaver, 1949), evenness (the equitability metric calculated in QIIME as: Shannon diversity / log2(number of observed OTUs)), and Balance-Weighted-abundance Phylogenetic Diversity (BWPD) (McCoy and Matsen IV, 2013). To assess the impact of variable sequencing depth on these diversity measures, rarefaction curves were made with multiple rarefactions from the lowest to the deepest sequencing depth, at intervals of 3000 sequences, with replacement and 100 iterations (Fig S1) using parallel_multiple_rarefactions.py, parallel_alpha_diversity.py and collate_alpha.py QIIME scripts. Alpha diversity metrics were then calculated using the mean of the 100 iterations of the deepest sequencing depth for each sample (McMurdie and Holmes, 2014). This approach was used to avoid losing data, and to estimate alpha diversity as accurately as possible. The Shannon index (OTU richness and evenness), and Equitability (evenness) were calculated using QIIME scripts as described above. The BWPD index that captures both the phylogeny (summed branch length) and the relative abundance of species was calculated using the guppy script with *fpd* subcommand (http://matsen.github.io/pplacer/generated_rst/guppy_fpd.html). Boxplots and statistical analyses were performed with IBM SPSS version 22.

To calculate the beta diversity between groups of samples (*e.g.* months or seasons), we used a non-rarefied OTU table to calculate two metrics that are robust to sequencing depth variation: weighted Unifrac (Lozupone et al,. 2007) and Jensen-Shannon divergence (JSD) (Fuglede and Topsoe 2004; Preheim *et al*., 2013). We used the phyloseq R package (version 1.19.1) (McMurdie and Holmes, 2013) (https://joey711.github.io/phyloseq) to first transform the OTU table into relative abundance (defined here as the counts of each OTU within a sample, divided by the total counts of all OTUs in that sample) then to calculate the square root of each metric (JSD or weighted UniFrac), and finally to perform principal coordinates analysis (PCoA) (Gower, 1966). As we observed potential arch effects with sqrt(JSD), we decided to use Nonmetric multidimensional scaling (NMDS, from the phyloseq package that incorporates the *metaMDS*() function from the R vegan package, Oksanen, J. *et al*., 2010. R package version 2.4-1) (Shepard, 1962; Kruskal, 1964) plots. A square root transformation is necessary here to transform weighted Unifrac (non Euclidean metric) and JSD (semi-metric) into Euclidean metrics (Legendre and Gallagher, 2001). Differences in community structure between groups (*e.g.* bloom vs. non-bloom samples) were tested using: (i) analysis of similarity (Clarke, 1993) using the *anosim*() function. The non-parametric Analysis of Similarity (ANOSIM; Clarke, 1993) has been used to test if the similarity among group sample is greater than within-group sample. If the *anosim*() function returns an R value of 1, this indicates that the groups do not share any members of the bacterial community. (ii) Differences in community structure between groups was also tested using permutational multivariate analysis of variance (PERMANOVA; Anderson, 2001) with the *adonis*() function. Both ANOSIM and PERMANOVA tests can be sensitive to dispersion, so we first tested for dispersion in the data by performing an analysis of multivariate homogeneity (PERMDISP, Anderson, 2006) with the permuted *betadisper*() function. In our analysis, we observed a significant dispersion effect when cyanobacterial sequences were included. The dispersion effect makes the PERMANOVA and ANOSIM results difficult to interpret. Dispersion mostly disappeared when we removed the cyanobacterial sequences, meaning that cyanobacteria were in part responsible for the differences in dispersion between groups. PERMANOVA, PERMDISP and ANOSIM were performed using the R vegan package (Oksanen, J. *et al*., 2010. R package version 2.4-1), with 999 permutations. Beta diversity analyses were also performed using a rarefied OTU table (rarefied to 10,000 reads per sample) and similar results were observed (data not shown). Phylogenetic trees used for phylogenetic analysis were built using FastTree (version 2.1.8, Price *et al*., 2009) (http://meta.microbesonline.org/fasttree). Three other tree inference methods were tested, yielding similar results to FastTree (Supplementary Methods).

### Bloom definition and K-means partitioning

Only a small subset of our samples were associated with estimates of cyanobacterial cell counts. We therefore estimated the relative abundance of cyanobacteria based on 16S rRNA gene amplicon data, which was significantly (but imperfectly) correlated with *in situ* cyanobacterial cell counts from a limited number of samples (Figure S6, adjusted R2=0.336; F1,50=27.46, P<0.001). The reason for the imperfect correlation is that, even when their absolute numbers are low, cyanobacteria can still dominate the community in relative terms.

To define cyanobacterial blooms, we followed the biological pulse disturbance definition described in Shade *et al*. (2012). Specifically, we defined a critical threshold of cyanobacterial relative abundance above which the Shannon diversity of the community begins to decline sharply, consistent with a major ecological disturbance (Figure S2). The decline in diversity is most pronounced when cyanobacteria make up 20% or more of the community, so we defined samples with 20% cyanobacteria or more as "bloom samples" (Table S7).

As an alternative and completely independent way of binning samples, we used the K-means partitioning algorithm (MacQueen, 1967), implemented with the function *cascadaKM*() from the vegan package in R, with 999 permutations. The OTU table was first transformed by Hellinger transformation (Rao, 1995) as advised in Legendre and Legendre (1998) by using the *decostand*(x, method=”hellinger”) function from R vegan package. OTU tables are generally composed of many zeros (as is the case for our data), which is inappropriate for the calculation of Euclidean distance. Hellinger transformation is a method to avoid this problem by down-weighting low-abundance OTUs (Legendre and Gallagher, 2001). We tested the partitioning of the 135 samples into 2 to 10 groups, based on the microbial community composition. The Calinski-Harabasz index (Caliński & Harabasz, 1974) was used to determine that our samples naturally clustered into two groups (Figure S3), and bloom samples (defined as above) were all found in a single K-means group (Figure S5). This suggests that the lake samples are naturally divided into two groups, and that cyanobacteria are a major distinguishing feature between groups.

### Changes in community composition over time

In order to investigate microbial community variation over time, we first analysed the change in Bray-Curtis dissimilarity over years. We performed separated analyses for littoral and pelagic OTU tables, after filtering out singleton OTUs only observed in one sample. This yielded 3491 OTUs for littoral samples and 3371 OTUs for pelagic samples. These two OTU tables were transformed to relative abundances prior to analysis. We calculated the Bray-Curtis dissimilarity between all pairs of samples using the QIIME script beta-diversity.py. We verified that distribution of Bray-Curtis dissimilarity across samples was approximately normal. Then, we used a custom script (see Supplementary file: “File_S2_R_scripts.txt”) to group the samples based on the amount of time (years) separating them, and to plot the mean dissimilarity of samples against their separation in time. Error bars were determined by calculating the standard error of the mean.

In a second approach, we used multivariate regression tree analyses (Breiman *et al*. 1984; De’ath 2002) with different time scales: year, season, month, week and day of the year. The goal here is to identify the temporal variables that best explain the variation in microbial community composition. An analysis was performed for each temporal variable (year, season, month or DoY) using the function *mvpart*() and *rpart.pca*() from the R mvpart package (Therneau and Atkinson, 1997; De'ath, 2007). Prior to analysis, the OTU table was Hellinger transformed (Rao, 1995) as advised in Ouellette *et al*., (2012). This approach is particularly useful to investigate both linear and non-linear relationships between community composition and a set of explanatory variables without requiring residual normality (Ouellette *et al*., 2012). After 100 cross-validations (Breiman *et al*. 1984), we plotted and pruned the tree using the 1-SE rule (Legendre & Legendre 2012) to select the least complex model, avoiding over-fitting. We then used the function *rpart.pca*() from mvpart package to plot a PCA of the MRT.

### Taxa-environment relationships

To investigate taxa-environment relationships, we performed a redundancy analysis (RDA; Rao, 1964) that searches for the linear combination of explanatory variables (the matrix of abiotic environmental data) that best explains the variation in a response matrix (the OTU table). The OTU table was transformed by Hellinger transformation (Rao, 1995) as advised in Legendre and Legendre (1998). The explanatory (environmental) matrix was first log-transformed then z-score standardized using the function *decostand*(x, method=”standardize”) because different environmental parameters are in different units. The environmental matrix variables included: total phosphorus in μg/L (TP), total nitrogen in mg/L (TN), particulate phosphorus in μg/L (PP, the difference between TP and DP), particulate nitrogen in mg/L (PN, the difference between TN and DN), soluble reactive phosphorus in μg/L (DP), dissolved nitrogen in mg/L (DN), 1-week-cumulative precipitation in mm, 1-week-average air temperature in Celsius and microcystin concentration in μg/L. The functions *corvif*(x) (Zuur *et al*., 2009) and *cor*(x, method=”pearson”) (the Pearson correlation; Bravais, 1846; Pearson, 1896) from the R stats package were applied to assess colinearity among explanatory variables (Table S2). Based on these correlation tests, we concluded that TP and TN were highly correlated with PP and PN, respectively, so TP and TN were removed. RDA was performed using the *rda*(scaling=2) function from the R vegan package. To determine the significance of constraints, we used the *anova.cca*() function from the R vegan package (Table S4A). Finally, we performed another RDA with all possible interactions between variable (except for Microcystin that is more a consequence of the bloom) to test if interactions between environmental variables could better explain the cyanobacterial bloom. The significance of the interactions is shown table S4B. Both RDAs were performed on a reduced dataset (a subset of 74 samples for which environmental data were available; see Supplementary file: File_S1_Environmental_Table.txt).

### Differential OTU abundance analysis

To identify genera and OTUs associated with blooms, we used the ALDEx2 R package (version: 1.5.0 (Fernandes *et al*., 2014). We used the aldex() function to perform a differential analysis with Welsh’s t-test and 128 Monte Carlo samples. ALDEx2 uses the centred log-ratio transformation to avoid compositionally issue. Taxa (OTUs or genera) with a Q-value below 0.05 after Benjamini-Hochberg correction were considered biomarkers. The top 25 biomarkers (with the highest differential scores) are listed in Table S8.

### Bloom classification

To classify bloom and non-bloom samples (Table S7), we used the Bayesian inference of microbial communities (BIOMICO) model described by Shafiei *et al*., (2015). This supervised machine learning approach infers how OTUs are combined into assemblages, and how combinations of these assemblages differ between bloom and non-bloom samples. An assemblage here is defined as a set of co-occurring OTUs. We defined bloom samples as described above, and trained the model with two different approaches: (i) with 2/3 of the total data, selected at random, and (ii) with two distinctive years: 2007, a year with only a short-lived fall bloom, and 2009, a year in which Fortin *et al*., (2015) observed a high biomass of cyanobacteria during the summer. In the training stage, BIOMICO learns how OTU assemblages contribute to community structure, and what assemblages tend to be present during blooms. In the testing stage, the model classifies the rest of the data (not used during training), and we assess accuracy as the percentage of correctly classified samples. To assess the performance of BIOMICO relative to a random classifier, we approximated a random classifier using a binomial distribution with correct classification probability of 0.5.

### Bloom prediction

We attempted to predict the timing of blooms using sequence or environmental data. As many OTUs or genera may have such low abundances that they might be missed in some samples, and might also increase the probability of finding spurious correlations, we pre-filtered the OTU table by removing taxa with summed relative abundances (over the 135 samples) lower than an arbitrary threshold of 0.1. Our goal was to predict the timing of the next bloom, using sequencing and/or environmental data from samples taken before a bloom event. Samples taken during a bloom were not used in these analyses. Thus, we used 21 samples with full environmental information when the analysis included these variables, and 54 samples when the analysis did not require the environmental variable. We defined the time (in days) from each non-bloom sample to the next bloom sample of the year as the dependent variable. In these analyses, we used either OTUs, genera, or environmental data, as predictor variables. We also calculated the trend in all predictor variables from one sample to the next by subtracting the latter values from the former and dividing by the number of days that separated the two sample dates. In this way, we obtained a trend value for each predictor variable.

Genetic programming, in the form of symbolic regression (SR) (Koza, 1992), is a particular derivation of genetic algorithms that searches the space of mathematical equations without any constraints on their form, hence providing the flexibility to represent complex systems, such as lake microbial communities. Contrary to traditional statistical techniques, symbolic regression searches for both the formal structure of equations and the fitted parameters simultaneously (Schmidt and Lipson 2009). There are however some caveats associated with SR. First, as with any other regression technique, overfitting may occur and measures that correct for model complexity, such as the Akaike information criterion (AIC,) should be used to compare equations. Second, contrary to standard regression techniques, there are no standard ways to interpret SR equations. Finally, SR suffers from the same limitations of evolutionary algorithms in general. In many cases the algorithm may get stuck in local minima of the search space, requiring time (or even a restart with different parameters) to find the global minimum. We used the software *Eureqa* (http://www.nutonian.com/products/eureqa/, version 1.24.0) to implement SR, using 75% of the data for model training and 25% for testing. As building blocks of the equations we used all predictor variables (including trends), random constants, algebraic operators (+, –, ÷, ×) and analytic function types (exponential, log and power). As no *a priori* assumptions regarding relationships between terms could be made, the search was fully unbounded. Given the inherent stochasticity of the process, ten replicate runs were conducted for each analysis. All runs were stopped when the percentage of convergence was 100, meaning that the formulas being tested were similar and were no longer evolving. Each run produces multiple formulas along a Pareto front (see Cardoso *et al*. 2015.). For each formula, we calculated the Akaike information criterion (AIC) and the corrected AIC (Burnham and Anderson, 2002) for small sample sizes. Based on *Eureqa* complexity (number of parameter) and Eureqa fit score (model accuracy), multiple formulae were selected from each of the ten runs (see Supplementary File: File_S3_SR_table.xlsx). The formula with the lowest AICc for each analysis was retained and considered the "best" formula (Table 2).

## Results

### Defining and characterizing blooms

To survey microbial diversity over time, we analysed 135 lake samples sequenced to an average depth of 54,231 reads per sample (minimum of 1000 reads per sample), and clustered the sequences into 4,061 operational taxonomic units (OTUs). Rarefaction curves showed that this depth of sequencing provided a thorough estimate of community diversity (Figure S1). To assess the repeatability and predictability of cyanobacterial blooms, we first needed to define bloom events. Instead of defining blooms based on cyanobacterial cell counts, we used a definition based on the extent to which the bloom disturbs the community. Above 20% cyanobacteria, Shannon diversity begins to decline sharply (Figure S2). We therefore used a 20% cutoff to bin our samples into "bloom" or "non-bloom" (Table S7).

Based on our definition, bloom samples necessarily have lower Shannon diversity than non-bloom samples (Figure 1). More surprisingly, bloom samples had significantly (Mann-Whitney test, U= 814, P<0.001) higher phylogenetic diversity (BWPD) compared to non-bloom samples (Figure 1A). These result suggests that cyanobacterial blooms lead to (i) an increase in phylogenetic diversity by adding additional, relatively long cyanobacterial branches to the phylogeny, and (ii) a decrease of taxonomic evenness due to the dominance of cyanobacteria. However, when we repeated the same analysis after removing all cyanobacterial OTUs, we found that blooms did not alter the diversity of the remaining (non-cyanobacterial) community (Figure 1 D, E, F). These exploratory alpha diversity analyses prompted us to investigate how community composition changed between bloom and non-bloom samples, and over time.

**Figure 1.**
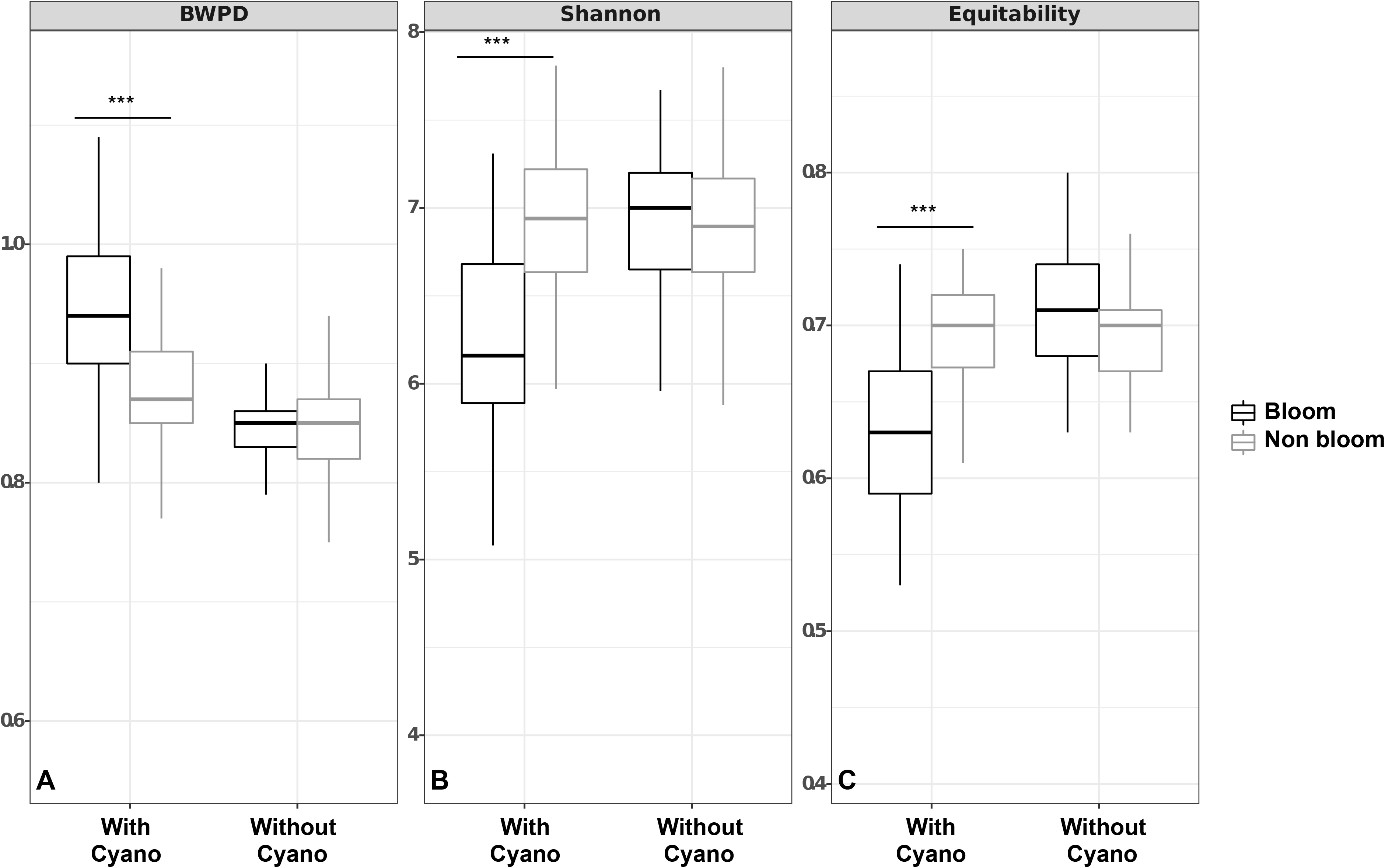
Comparison of alpha diversity between bloom and non-bloom states. Three alpha diversity metrics were employed: (A) BWPD, (B) the Shannon index, and (C) the Shannon evenness (equitability) to compare alpha diversity between bloom (black) and non-bloom (grey) samples. We repeated the same analysis after removing Cyanobacteria. Comparisons were performed using a Mann-Whitney test (* P < 0.05, ** P < 0.01, *** P < 0.001).

Despite their limited impact on the diversity of the non-cyanobacterial community, we found that blooms clearly alter the community composition of the lake. Using weighted UniFrac distances to assess differences in community composition, we observed a separate grouping of bloom and non-bloom samples (Figure 2A). However, the difference in community composition could not be assessed with PERMANOVA statistics because bloom and non-bloom samples were differently dispersed (Table S6. When we removed the Cyanobacteria counts and re-normalized the OTU table (Figure 2B), we still observed a significant, but less pronounced difference between bloom and non-bloom samples (PERMANOVA, R2=0.035; P<0.001; ANOSIM R=0.211; P<0.01; PERMDISP P = 0.084; Table S6). We observed the same trend using another beta diversity metric, JSD (Table S6 and Figure S7). These results suggest that even excluding Cyanobacteria (the bloom-defining feature), the bloom community still differs to some extent from the non-bloom community.

**Figure 2.**
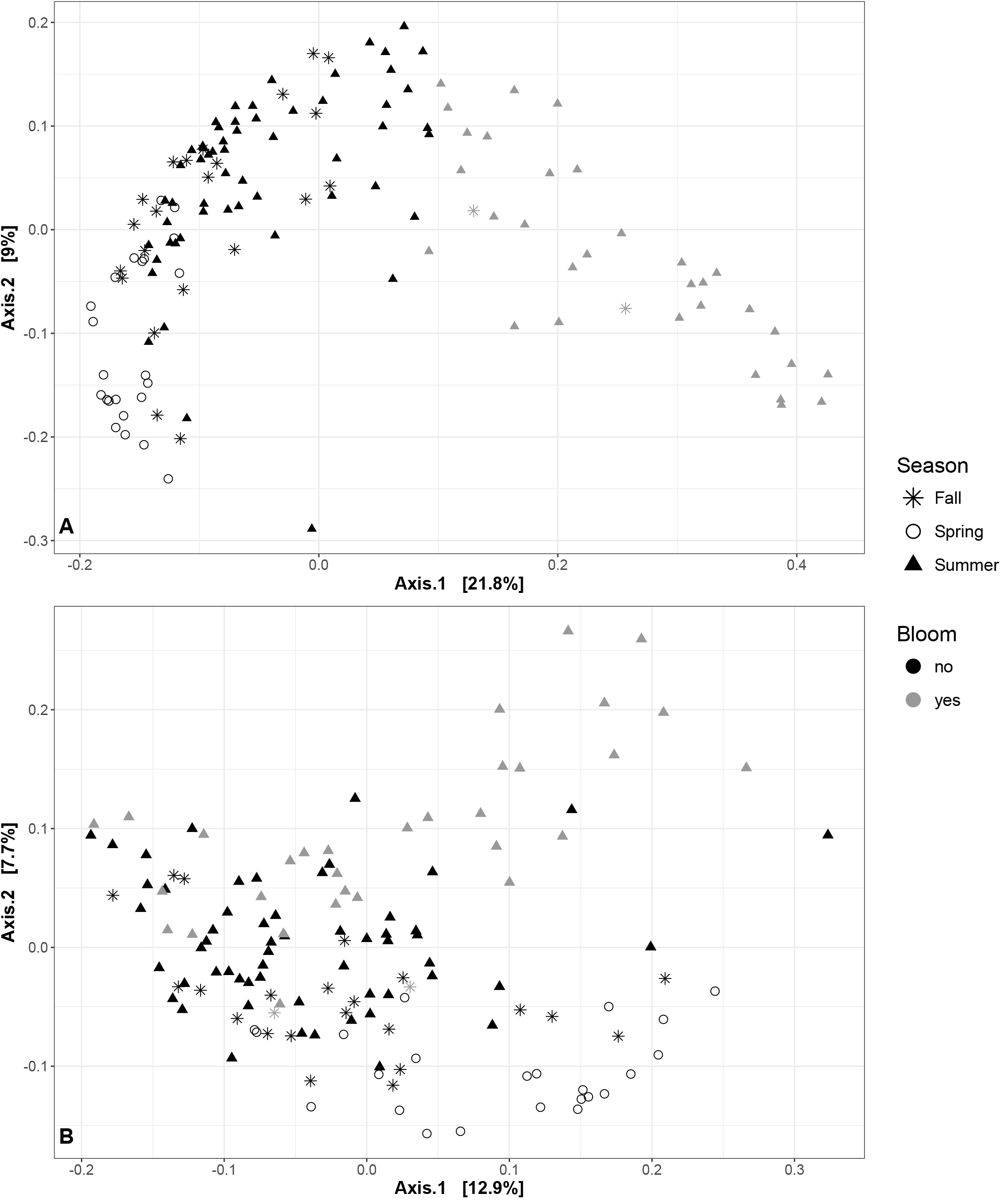
Changes in community composition across seasons and bloom events. Each point in the PCoA plot represents a sample, with distances between samples calculated using weighted UniFrac as a measure of community composition. Non-bloom samples are shown in black, bloom samples in grey. Different shapes describe the different seasons: circle for Spring, triangle for Summer and star for Fall. (A) Samples with all OTUs included. (B) Samples excluding OTUs from the phylum Cyanobacteria.

### Abiotic factors associated with blooms

A subset of our samples was associated with environmental measurements that might explain bloom events. We performed an RDA to identify environmental variables that could explain how bloom and non-bloom samples are grouped, and found particulate nitrogen (PN), particulate phosphorus (PP), microcystin concentration, and to a lesser extent soluble reactive phosphorus (DP), to be most explanatory of the bloom (Figure S8; adjusted R2=0.273; ANOVA, F7,66=4.919, P<0.001). DN and temperature explain less variation and act in opposing directions (Pearson correlation = -0.18), perhaps because higher temperatures favour the growth of microbes that rapidly consume dissolved nitrogen (Hong *et al*., 2014). Together, these environmental variables explain ~25% of the microbial community variation (axis 1: 18.5%; axis 2: 6.9%) suggesting that unmeasured biotic or abiotic factors are needed to explain the remaining ~75% of the variation. We also explored the ability of interactions among environmental variables to explain variation, but despite the modest increase in R2 to 0.34 (to be expected given the added variables) we did not observe any significant interactions (Supplementary Table 4B).

### Community dynamics vary more within than between years

We next asked how the lake microbial community varied over time, at scales ranging from days to years. As described above, samples can be partially separated according to season (spring, summer or fall) based on weighted UniFrac distances (Figure 2). However, seasons differed significantly in their dispersion (with summer samples visibly more dispersed in Figure 2), violating an assumption of PERMANOVA and ANOSIM tests, and preventing us from determining whether samples varied more by months, seasons or years (Table S6). However, it is visually clear from Figure 2 that bloom samples explain much of the variation in summer community composition.

To more clearly track changes in community composition over time (temporal beta diversity), we calculated the Bray-Curtis dissimilarity between pairs of samples separated by increasing numbers of years. We did not observe any tendency for the community to become more dissimilar over time, suggesting a long-term stability of the bacterial community on the time scale of years in both the littoral (linear regression, F(1,1999)=1.171, P>0.05) and pelagic sampling sites (linear regression, F(1,2078)=0.8467, P>0.05; Figure S4). Consistently, even though years differed significantly in their dispersion (PERMDISP *P* < 0.05), community composition remained relatively similar from year to year. (Weighted Unifrac: ANOSIM R<0.1, P<0.010; PERMANOVA R^2^=0.011, P=0.098).

To further explore temporal signals in the data, we used a multivariate regression tree (MRT) approach to determine how community structure varies over time scales of days to years. Consistent with the stable Bray-Curtis similarity over years (Figure S4), we found that year-to-year variation explains very little of the variation in community structure (R^2^=0.027; Table S5). Week of the year explained the most the community variation (R^2^=0.274; Figure 3, Table S5), followed closely by day (R^2^=0.254; Table S5) and month (R^2^=0.216; Table S5). Even though weeks explained the most variation, much of this variation is captured at longer time scale of months. Figure 3 shows how the regression tree roughly divides samples by season: Split 1 (red) corresponds to samples taken before May 12 (early spring), split 2 (green) to samples taken between May 12 and June 23 (late spring), split 3 (yellow) to samples taken after October 6 (fall), split 4 (blue) to samples taken between June 23 and July 14 (early summer), split 5 (cyan) to samples taken between July 14 and August 11 (mid-summer), and split 6 (purple) to samples taken between August 11 and October 6 (late summer). The PCA ordination based on MRT (Figure 3B) shows that community dynamics appear to be somewhat cyclical, returning to roughly the same composition each year. Different times of year are characterized by different sets of OTUs, for example AcI-B1 and PnecB in early summer and *Microcystis* and *Dolichospermum* in mid-summer.

**Figure 3.**
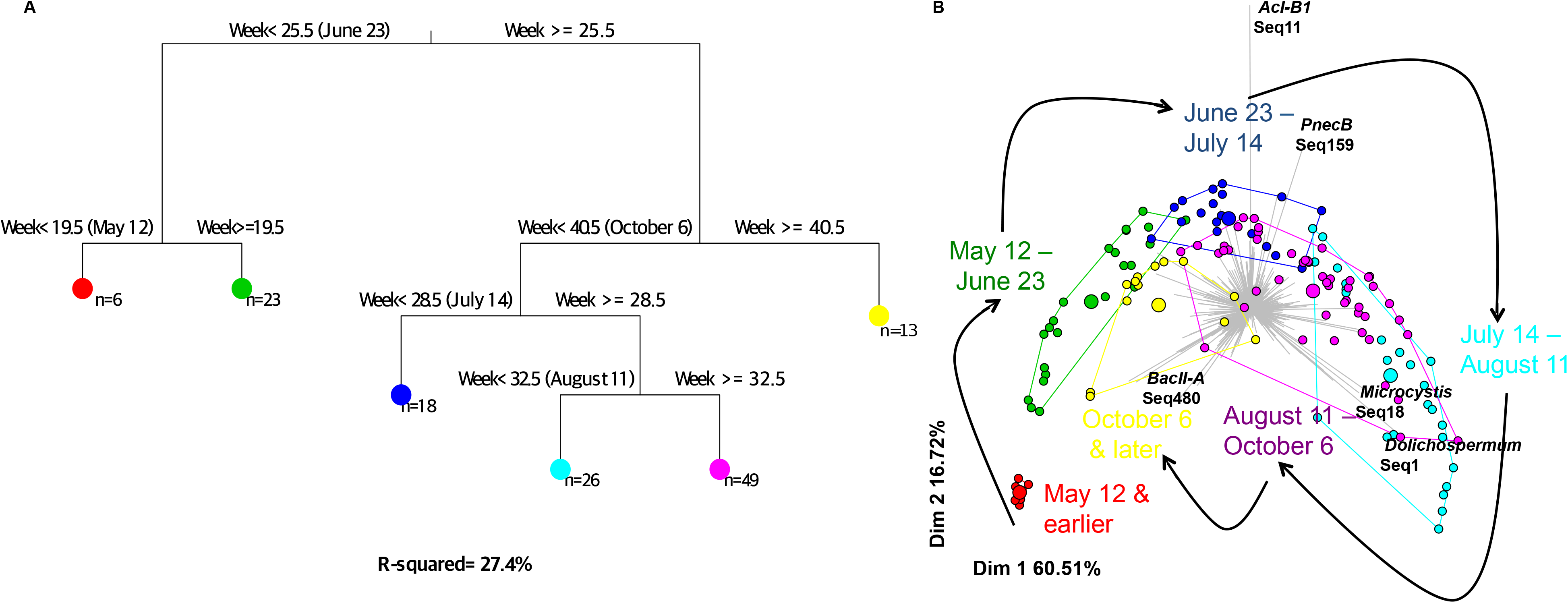
Cyclical community composition dynamics. Multivariate regression tree (MRT) analysis was used to estimate the impact of time on bacterial community structure. (A) The most parsimonious tree shows how the community is partitioned by MRT using week of the year as a temporal variable. Six different leaves (large coloured circles) were defined based on microbial abundance and composition. (B) The community composition within leaves is represented in a PCA plot, where small points represent individual samples and large points represent the group mean (within the leaf). The grey barplot in the background indicates OTUs whose differential abundance explains variation in the PCA plot.

To determine if the variation observed during summer (Figures 2 and 3) could be driven by cyanobacterial bloom events, we repeated the MRT analyses after removing all cyanobacterial sequences. Similar MRT results were obtained after removing cyanobacteria, suggesting that the entire bacterial community, not just cyanobacteria, are responsible for temporal variation (Table S5). Together, these results show how bacterial community dynamics follow an annually repeating, cyclical pattern, and that both cyanobacteria and other bacteria contribute to the dynamics.

### Blooms are repeatably dominated by Microcystis and Dolichospermum

To explore potential biological factors involved in bloom formation, we attempted to identify taxonomic biomarkers of bloom or non-bloom samples, at the genus and OTU levels. To do so, we performed a differential analysis using ALDEx2 to identify the genera or OTUs that are most enriched in bloom samples. We found several significant biomarkers and as expected, the strongest bloom biomarkers belonged to the phylum Cyanobacteria (Table S8). The two strongest OTU- and genus-level biomarkers were *Microcystis* (Microcystacae) and *Dolichospermum* (Nostocaceae, previously named *Anabaena*), both genera of Cyanobacteria.

### Blooms can be accurately classified based on non-cyanobacterial sequence data

Given the observation that bloom samples have distinct cyanobacterial and non-cyanobacterial communities (Figure 2), we hypothesized that blooms could be classified based on their bacterial community composition. We trained a machine-learning model (BIOMICO) on a portion of the samples, and tested its accuracy in classifying the remaining samples (Methods). BIOMICO was able to correctly classify samples with ~92% accuracy (Table 1). Such high accuracy is expected because blooms are defined as having >20% cyanobacteria, so the model should be able to easily classify samples based on cyanobacterial abundance.

**Table 1.** Bloom classification results. We used a supervised machine learning approach (BioMico) to determine if samples can be classified into bloom bins based on microbial assemblages (Methods). Accuracy was calculated as the percentage of correctly classified samples (true positives + true negatives) relative to the total number of samples in the testing set The 95% confidence intervals of a random classifier (Methods) and the *P*-values (that the real classifier differs from random) are also shown.

| **Training set** | **Testing set** | **Classification Accuracy** | **False positives** | **False negatives** | **True negatives** | **True positives** | **95% confidence interval of random classifier** | **P-value (real classifier differs from random)** |
| --- | --- | --- | --- | --- | --- | --- | --- | --- |
| 2/3 of all samples | 1/3 of all samples | 91.84 % | 4 | 0 | 33 | 12 | 36-64% | $8.225 \times 10^{-10}$ |
| 2007 & 2009 samples | All other samples | 92.52% | 8 | 0 | 73 | 26 | 40-60% | $< 2.2 \times 10^{-16}$ |
| 2/3 of all samples, without cyanobacteria | 1/3 of all samples, without cyanobacteria | 85.71% | 6 | 1 | 31 | 11 | 36-64% | $3.625 \times 10^{-07}$ |
| 2007 & 2009 samples, without cyanobacteria | All other samples, without cyanobacteria | 83.18% | 9 | 9 | 72 | 17 | 40-60% | $1.781 \times 10^{-12}$ |

In a more challenging classification task, BIOMICO was able to classify samples with 83-86% accuracy after excluding cyanobacterial sequences. This result supports the existence of a characteristic non-cyanobacterial community repeatably associated with the bloom. Two different training approaches (Methods) yielded similar classification accuracy, both significantly better than random (Table 1), but found different bloom-associated assemblages. When we compared the best assemblages obtained with the two different trainings, focusing only on the 50 best OTU scores, only 11 OTUs were found in both trainings (Table S9). This result suggests that data can be classified into bloom or non-bloom samples, but different assemblages (containing different sets of OTUs) can be found with similarly high classification accuracy (Table S9). This is consistent with a general lack of repeatability at the level of individual OTUs, but that there exist combinations of OTUs (Table S8) that are characteristic of blooms.

### Blooms can be predicted by sequence data

The existence of microbial taxa and assemblages characteristic of blooms suggests that blooms could, in principle, be predicted based on amplicon sequence data. We therefore used symbolic regression (SR) to model the response variable “days until bloom” as a function of OTU- or genus-level relative abundances, their interactions, and their trends over time (Methods). To achieve true prediction, not simply classification, we used only samples collected prior to each bloom event in order to predict the number of days until a bloom sample (*i.e.* bloom samples themselves were not used). We based our analysis on 54 samples, ranging from 7 to 112 days before a bloom sample. Due to limitations in the resolution of sampling (approximately weekly), we cannot know the exact start date of a bloom, only the first date sampled. Using OTUs or genera, we were able to predict the timing of the next bloom event with 80.5% or 78.2% accuracy on tested data, respectively (Table 2). Using a subset of 21 samples with a full complement of environmental data, we were able to compare the predictive power of sequence data (OTU or genus level) versus environmental data. Predictions based on genus-level sequence data clearly outperformed predictions based on environmental data. Predictions based on OTU-level sequence data explained less variance than predictions based on genera, consistent with OTUs being more variable and less reliable bloom predictors than higher taxonomic units.

**Table 2.**
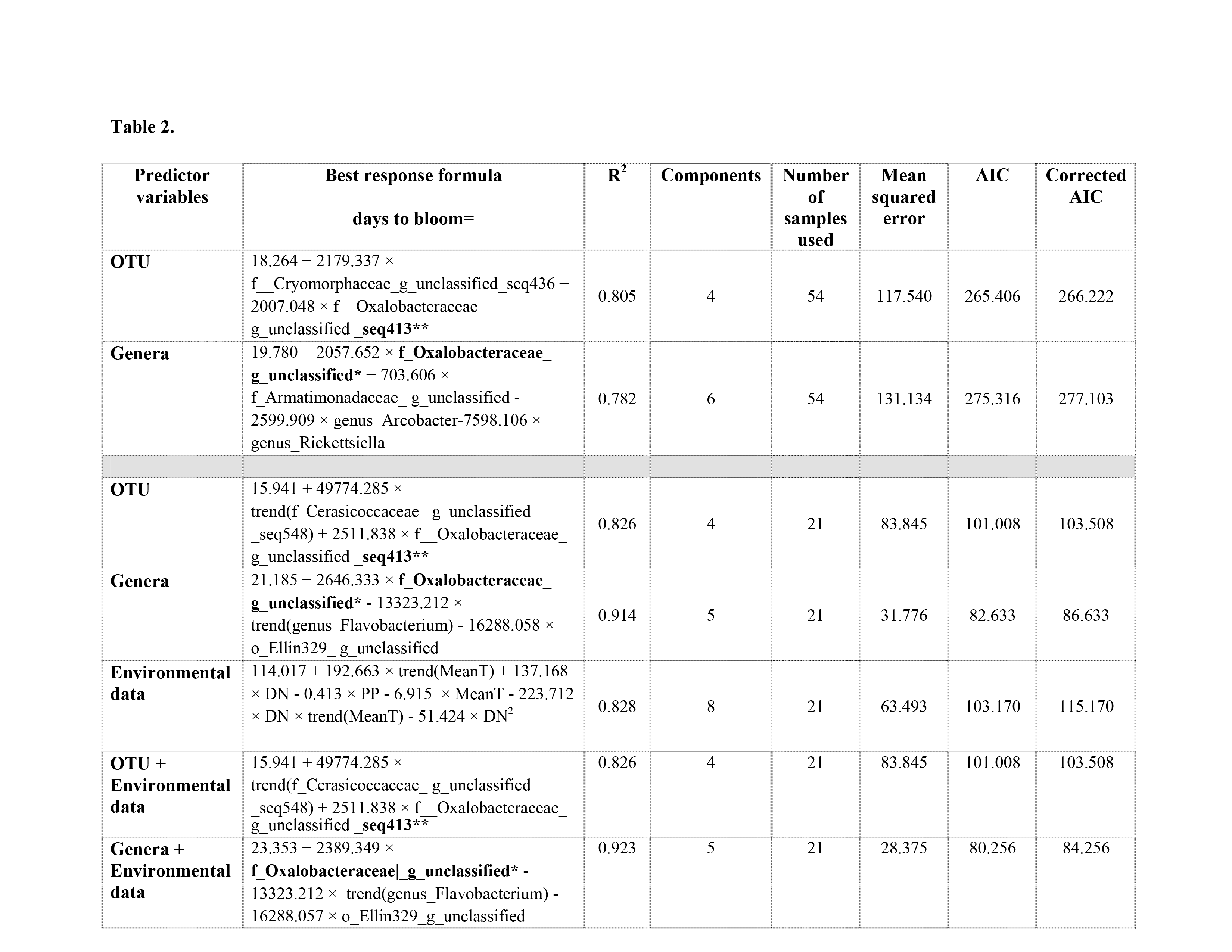
Predicting bloom timing with symbolic regression (SR). The best formula found by SR is shown for each category of predictor variables. SR was performed on two datasets. First, OTUs and genera were used as predictor variables, using the maximum number of non-bloom samples (*N =* 54). Second, in order to determine the impact of including environmental data as predictor variables, we used only samples with a full set of metadata (*N* = 21). (*/** indicate OTUs/genera found multiple times in SR formulas).

All models tend to overshoot when based on samples taken closer to the bloom (*i.e*. negative residuals), and tend to predict bloom events too soon when based on samples farther from the bloom (Figure S9). One taxon – a member of the order Burkholderiales in the family Oxalobacteraceae (unknown genus; Greengenes taxonomy) was consistently found in every predictive formula (Table 2). At the OTU level, seq413 (Table 2) is assigned to Oxalobacteraceae by Greengenes (with 67% confidence) but to *Polynucleobacter* C-subcluster (with 99% confidence) based on TaxAss, a freshwater-specific database (Table S10). While *Microcystis* and *Dolichospermum* are dominant closer to bloom events, seq413 showed the opposite pattern, decreasing in relative abundance as the bloom approaches (Figure 4). The fact that seq413, but not *Microcystis* or *Dolichospermum*, appears in the predictive equations suggests that the decline in Oxalobacteraceae/seq413 is detectable before the increase in Cyanobacteria. Indeed, seq413 appear to decline before *Microcystis* or *Dolichospermum* increase (Figure 4). However, the predictive analyses were done at the OTU or genus level, such that Cyanobacteria were not treated as one entity (*i.e.* one variable in predictive equations). It is therefore possible that the decline in seq413 was driven by a total increase in the sum of all Cyanobacteria, none of which could be detected individually. To test this possibility, we repeated the SR analysis after merging Cyanobacteria into a single variable, and found that Cyanobacteria were never found in any predictive equation. This is consistent with Oxalobacteraceae/PnecC declining before Cyanobacteria increase. Hence, changes in the microbial community provide information about impending blooms before they occur.

**Figure 4.**
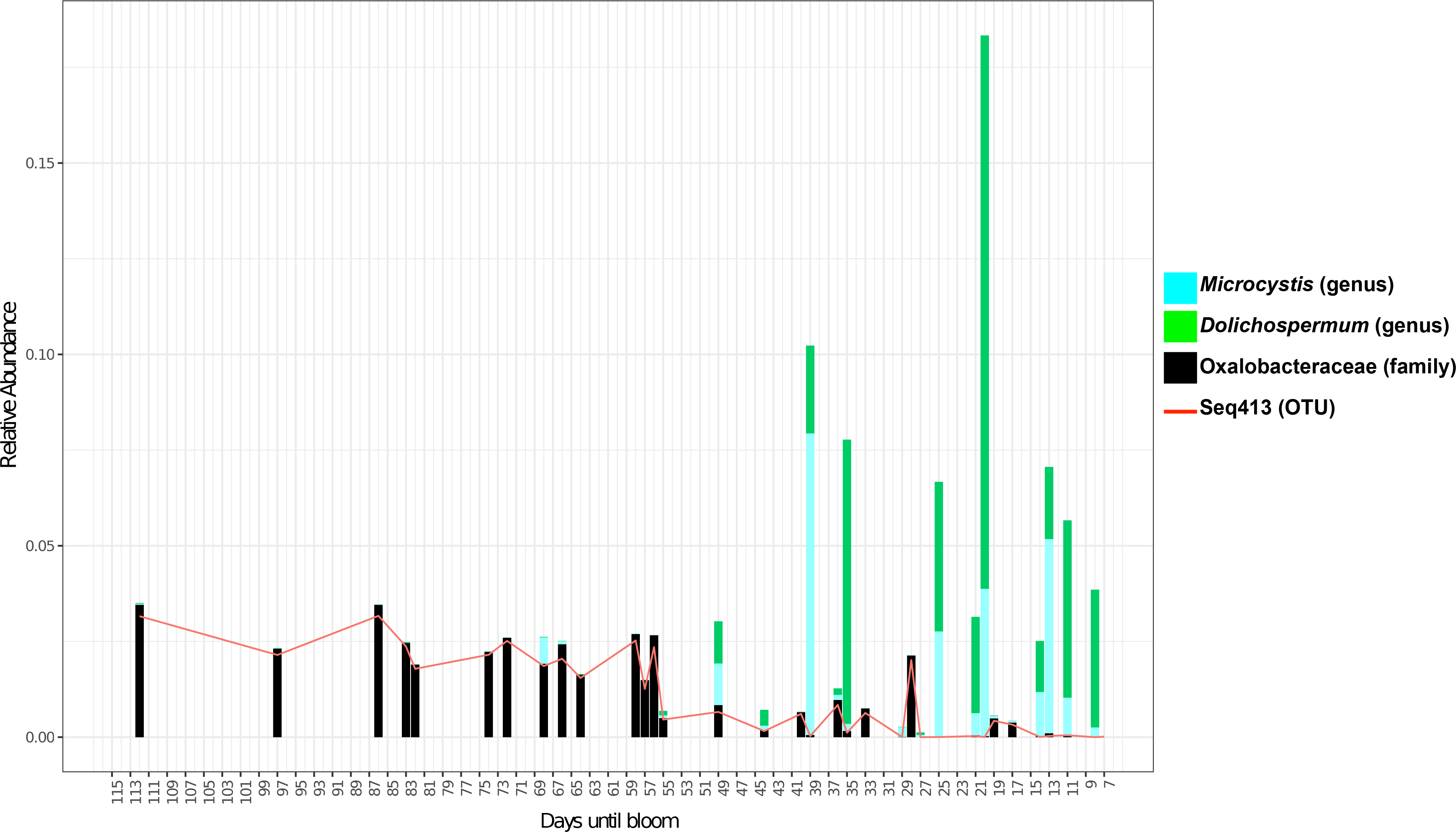
Oxalobacteraceae and seq413 decline while *Microcystis* and *Dolichospermum* increase as a bloom event approaches. We plotted the relative abundance of relevant taxa from 112 to 7 days before a bloom sample. Oxalobacteraceae (genus unclassified) and the OTU seq413 (Oxalobacteraceae, genus unclassified or *Polynucleobacter* PnecC) are relatively abundant long before a bloom event, and gradually decline as bloom events approach. *Microcystis* and *Dolichospermum* are the two most dominant bloom-forming cyanobacteria.

## Discussion

We used a deep 16S rRNA amplicon sequencing approach to profile the bacterial community in Lake Champlain over eight years, spanning multiple cyanobacterial blooms. We sequenced with sufficient depth that bacterial diversity estimates reached a plateau (Figure S1), and proposed a bloom definition based upon cyanobacterial relative abundance in 16S data. Although there is no consensus bloom definition, the World Health Organization has proposed guidelines, based on cyanobacterial cell density, to connect blooms to potential health risks (WHO, Guidelines for safe recreational water environments, 2003). We found that, while cyanobacterial relative abundance in 16S data is significantly correlated with cyanobacterial cell density, the correlation is imperfect (Figure S6) because cyanobacteria can have high relative abundance without achieving a high absolute cell density. Our bloom definition, based on relative, not absolute abundance is therefore more a measure of how cyanobacteria impact their surrounding bacterial community than a direct measure of human health risks.

Our results should be interpreted in light of four methodological caveats. First, the OTU data are compositional, such that only the relative OTU abundances are meaningful, and the relative abundances are non-independent (Gloor and Reid, 2016). As a result, removing certain OTUs or taxa (*e.g.* Cyanobacteria, as discussed in the paragraph below) does not remove their influence on the rest of the data. For some purposes, corrections for compositionality can be performed (*e.g.* ALDEx performs a centered log transform before inferring differentially abundant OTUs). BioMico might identify OTUs that are not truly associated with blooms, but that are falsely correlated with OTUs that are truly associated. However, this is not a major problem because the goal of BioMico is bloom classification, not identification of bloom-associated OTUs. A similar logic applies to prediction with SR: if the goal is pragmatic prediction, whether the predictive taxa are biologically meaningful (or mere artefacts of compositionality) is irrelevant. In reality, the fact that SR repeatably converged on equations with the same taxa (Table 2) suggests that these taxa are indeed biologically meaningful. The second caveat is that the same data was used to define blooms and also to classify/predict blooms, which could be considered circular reasoning. However, the bloom definition was based on a univariate summary of the data (Shannon diversity), while BioMico classification uses the multivariate data (the relative abundance of each OTU across samples). Therefore, circularity is limited because blooms were defined based on one feature of the data (a decline in Shannon diversity), and classification was based on a different feature (OTU identities). For the prediction task, circularity was limited because only non-bloom samples were used to predict the timing of a bloom event. The third caveat is that phylogenetic measures of alpha and beta diversity (BWPD and UniFrac, respectively) rely on a phylogenetic tree, which may be inaccurate. However, trees inferred using FastTree, ML or neighbour-joining gave very similar results (Supplementary Methods), so we expect tree errors to have a limited impact on our conclusions. The fourth caveat is that the choice of OTU calling will influence the number and identify of OTUs. We used a distribution-based OTU caller (Preheim *et al*., 2013), which uses the distribution of OTUs across samples to reduce the number of false-positive OTUs (*e.g.* due to sequence errors). Other methods, such as DADA2 (Callahan *et al*., 2016), oligotyping or minimum entropy decomposition (Eren *et al*., 2013; 2015), are similarly able to de-noise 16S data, while calling OTUs at fine taxonomic resolution (*e.g.* 99% rather than 97% identity). In the future, these methods could be used to analyze bloom dynamics at finer taxonomic resolution than the 97% cutoff used here.

Our results suggest that blooms decrease community diversity because of an increase in the relative abundance of cyanobacteria, not due to a reduction in the diversity of other bacteria. This result is based on an analysis of three diversity measures, before and after removing cyanobacterial sequences (Figure 1). Before removing Cyanobacteria, bloom samples clearly have lower Shannon diversity and evenness compared to non-bloom samples (this is true by definition, based on the nature of our bloom definition). After removing Cyanobacteria, there is no apparent difference in diversity or evenness. Removing cyanobacterial reads does not remove their influence on other OTUs, because of the dependence structure of compositional data (Gloor and Reid, 2016; Morton *et al*., 2017). However, even if removing Cyanobacteria creates a bias in the rest of the data, the same bias is introduced in both bloom and non-bloom samples alike, so the comparison should remain valid. The removal of cyanobacterial reads is analogous to the common practice of first removing eukaryotic reads from 16S data, and continuing all subsequent analyses on bacterial reads only. The dataset as a whole is biased by the removal of eukaryotes (*i.e.* the data becomes a 'subcomposition') but all samples have the same bias, so it is still possible to compare among samples. Regardless, these diversity comparisons (Figure 1) were exploratory in nature, and served as an entry point for more detailed beta diversity analyses, classification, and prediction.

Consistent with our current knowledge of temperate lakes (Shade *et al*., 2007; Crump *et al*., 2005), we found that community structure varied more within years than between years (Figures 2, 3, and S4; Tables S5 and S6). In agreement with previous observations in eutrophic lakes (Shade *et al*., 2007), Lake Champlain appears to return to a steady-state (Figure S4, Table S5), despite the biological disturbance induced by dramatic bloom events. Various studies have already shown temporal patterns in microbial community structure (Hofle *et al*., 1999; Lindstrom *et al*., 2000; Crump et al., 2003; Shade *et al*., 2007; Kara et al., 2013; Fuhrman *et al*., 2015), but ours does so in the context of cyanobacterial blooms.

The RDA results (Figure S8) are consistent with many previous studies describing the environmental factors responsible for blooms (Owens and Esaias, 1976; Hecky and Kilham, 1988). For example, cyanobacterial growth is optimal at higher temperatures, between 15 and 30oC (Konoka and Brock, 1978). We confirmed that cyanobacterial blooms are correlated with, and likely respond to nutrient concentrations, as previously described (Fogg, 1969; Jacoby *et al*., 2000; Paerl and Huisman, 2008; Paerl and Huisman, 2009, Fortin *et al* 2015, Isles *et al*., 2015). Dissolved nitrogen and temperature were negatively correlated, which could be explained by the fact that the lake becomes enriched in nitrates during spring, when temperatures are lower, and rain and drainage bring nutrients into the lake (Shade *et al*., 2007; Fortin *et al*., 2015). Another explanation would be that in the spring, before most of the bloom events occur, the majority of the nitrogen is dissolved, but when cyanobacteria and other phytoplankton increase in abundance over the summer, nitrogen becomes concentrated in particulate forms within cells. We found that measured abiotic variables explained only a part (~25%) of the variation between bloom and non-bloom samples. Including interactions between variables in the model increased the adjusted R^2^ to ~35%; however no significant interactions were found (Table S4B). The rest of the variation could be explained by unmeasured variables, such as different nitrogen species, water column stability and mixing (although Missisquoi Bay is shallow [~2-5m] and likely never stratified), or time-lagged variables. More variance might also be explained with a larger dataset containing more samples.

In addition to environmental variables, we showed that biological variables, in the form of bacterial OTUs or genera, also characterize bloom events. Differential analysis using ALDEx2 identified *Microcystis* and *Dolichospermum* as the top bloom biomarkers (Table S8). These two bloom-forming genera are associated with lake eutrophication (O’Neil *et al*., 2012) and are also known to produce cyanotoxins (Gorham and Carmichael *et al*., 1979; Carmichael, 1981). We found additional bloom biomarkers in the genus *Pseudanabaena* and the family Cytophagacaea, previously found to be associated with cyanobacterial blooms (Rashidan and Bird, 2001; O’Neil *et al*., 2012). The order Chthoniobacterales (in the phylum Verrucomicrobia) was also found as a bloom biomarker, consistent with previous studies that observed this taxon in association with *Anabaena* blooms (Louati *et al*., 2015). Other studies have reported specific association between Verrucomicrobia and Cyanobacteria, suggesting that members of this phylum might assimilate cyanobacterial metabolites (Parveen *et al*., 2013; Louati *et al*., 2015). We also found N2–fixing members of *Rhizobiales* order as bloom biomarkers. These taxa might be associated with the non-N2-fixing cyanobacteria *Microcystis*, potentially supporting its growth.

Using machine learning, we were able to classify bloom samples with high accuracy based on microbial assemblages, confirming that there is a specific microbial community associated with blooms. Consistent with the ALDEx2 results, *Microcystis* and *Dolichospermum* were present in all bloom assemblages (Table S9). Cyanobacterial blooms have been previously suggested to alter the local environment and the surrounding microbial community (Louati *et al*., 2015). As a result, these assemblages may include bacteria that are reliant on cyanobacterial metabolites and biomass. For example, we found that bloom assemblages included potential cyanobacterial predators from the order Cytophagales and the genus Flavobacterium (Table S9), both associated with bloom termination (Rashidan and Bird, 2001; Kirchman, 2002) but also taxa such as Methylophilaceae, acI, and acIV that have been previously associated with cyanobacterial blooms (Li *et al*., 2015; Woodhouse *et al*., 2016). We found that acI was abundant in early summer, just before the *Microcystis* and *Dolichospermum* blooms of mid-summer (Figure 3B). While acI might help "set the stage" for a bloom, acIV might have the capacity to use metabolites from cyanobacterial decomposition, and Methylophilaceae is a potential microcystin degrader (Bogard *et al*., 2014; Ghylin *et al*., 2014, Mou *et al*., 2013).

Finally, we show the potential for bloom events to be predicted based on amplicon sequence data. We acknowledge that long-term environmental processes such as global warming, and punctual seasonal events such as floods and droughts, are major determinants of whether a bloom will occur in a given year (Paerl and Huisman, 2008; Paerl and Paul, 2012). For example, no bloom occurred in 2007, likely due to a spring drought which dramatically reduced nutrient run-off into the lake. However, sequence data might be useful to predict bloom dynamics on shorter time scales of days, weeks or months. We demonstrated that it is possible to use pre-bloom sequence data to predict the number of days until a bloom event, with errors on the order of weeks (Figure S9) – the best that could be expected, given that sampling density was also on the order of weeks. Sequence data appears to be a strong predictor, similar or better than prediction with environmental variables (Table 2). These results are consistent with a recent study suggesting that abiotic environmental factors could be crucial to initiate blooms, but that biotic interactions might also be important in the exact timing and dominant members of the bloom (Needham and Fuhrman, 2016). Similarly, environmental variables explained relatively little variation in freshwater bacterial composition, while biotic variables (*i.e.* phytoplankton) explained more (Kent *et al*. 2004). It is possible that measuring more environmental variables, or using more complex time-lagged environmental variables (beyond the simple trends used in SR equations) could provide better predictions. However, microbial variables (OTUs) can be measured nearly exhaustively in a single sequencing run, whereas it is hard to know which environmental variables to measure (*e.g.* temperature, pH, nitrogen, etc. seem relevant but what about Fe, As, Mg, etc.?) and hard to measure them all in high-throughput. However, SR models might be prone to overfitting, which might explain why better predictive accuracy is achieved with fewer samples (Table 2). Our samples were rarely taken more often than weekly, explaining why prediction error is on the order of weeks (Figure S9). We expect that more samples taken over shorter time periods will reduce both overfitting and prediction error. We also note that the "best" predictive equations found by SR are not necessarily global optima, because the space of possible equations is not explored exhaustively.

Surprisingly, we never found Cyanobacteria as a bloom predictor in any of the predictive models (Table 2). This means that the models are not simply tracking a positive trend in cyanobacterial abundance, possibly because bloom events are "spiky" (Figure 4) and hence difficult to predict with weekly sampling. Instead, predictive equations always included a member of the order Burkholderiales, classified as Oxalobacteraceae with 67% confidence by Greengenes, or *Polynucleobacter* C (PnecC) with 99% confidence by TaxAss. We acknowledge this taxonomic uncertainty, but give preference to the higher-confidence PnecC assignment. PneC tends to be relatively abundant further ahead of bloom events (Figure 4). This observation could be explained by an ecological succession between PnecC and *Microcystis*/*Dolichospermum*. The fact that PnecC was chosen as a better predictor than Cyanobacteria suggests that PnecC begins to decline before any detectable increase in Cyanobacteria, providing a potential early warning sign. Šimek *et al*., (2011) showed that some PnecC taxa grow poorly in co-culture with algae, suggesting that negative interactions could also occur with cyanobacteria.

We have shown that cyanobacterial blooms contain highly (but not exactly) repeatable communities of Cyanobacteria and other bacteria. It appears that the community begins to change before a full-blown bloom, suggesting that sequence-based surveys could provide useful early warning signals. While the predictions of our models are fairly coarse-grained (*e.g.* prediction error on the order of weeks), they suggest that more accurate prediction might be enabled with increased sampling frequency. It remains to be seen to what extent bloom and pre-bloom communities – which show repeatable dynamics within one lake – are also repeatable across different lakes, and to what extent predictors could be universal or lake-specific. To improve predictions going forward, we suggest sampling additional lakes with dense time-courses, paired with 16S or metagenomic sequencing. In order to predict not just blooms but also the toxicity of blooms, sequencing should be paired with detailed toxin analyses.

## Data availability

Raw sequence data have been deposited NCBI GenBank under BioProject number PRJNA353865.

## Author information

The authors declare no competing financial interest.

## Acknowledgments

We thank Joe Bielawski, Lawrence David, Yonatan Friedman, Catherine Girard, Alan Hutchison, Jean-Baptiste Leducq, Pierre Legendre, Julie Marleau, Simone Perinet, Sarah Preheim, Zofia Taranu, Justin Silverman, Gavin Simpson and Amy Willis for advice, help in the laboratory and/or with data analysis. We thank three anonymous peer reviewers for their detailed and constructive suggestions. We also thank everyone who participated in sampling, data collection and analysis, with special thanks to David Juck, Alberto Mazza and Miria Elias. This research was funded by a Natural Sciences and Engineering Research Council (NSERC) Discovery grant and a Fonds de Recherche du Québec Nature et Technologies (FRQNT) New Researcher grant to BJS, and the federal government interdepartmental Genomics Research and Development Initiative (GRDI). NT is funded by a project from the European Union’s Horizon 2020 research and innovation program under the Marie Sklodowska-Curie grant agreement No 656647.

